# Regional differences in gene regulation may underlie patterns of sensitivity to novel insecticides in Colorado potato beetle

**DOI:** 10.1101/794446

**Authors:** Galen P. Dively, Michael S. Crossley, Sean D. Schoville, Nathalie Steinhauer, David J. Hawthorne

## Abstract

Agricultural insect pests frequently exhibit geographic variation in levels of insecticide resistance, which are often presumed to be due to the intensity of insecticide use for pest management. However, regional differences in the evolution of resistance to novel insecticides suggests that other factors are influencing rates of adaptation. We examined LC_50_ bioassay data spanning 15 years and six insecticides (abamectin, imidacloprid, spinosad, cyantraniliprole, chlorantraniliprole, and metaflumizone) for evidence of regional differences in Colorado potato beetle (CPB) baseline sensitivity to insecticides as they became commercially available. We consistently found that CPB populations from the Western USA had the highest baseline sensitivity to novel insecticides, while populations from the Eastern USA had the lowest. Comparisons of gene expression between populations from these regions revealed constitutively elevated expression of an array of detoxification genes in the East, but no evidence of additional induction when exposed to imidacloprid. Our results suggest a mechanism for geographic variation in rates of adaptation to insecticides whereby baseline levels of gene expression determine a population’s response to novel insecticides. These findings have implications for the regional development of insecticide resistance management strategies and for the fundamental question of what determines the rate of adaptation to insecticides.

## Introduction

Colorado potato beetle (CPB) is a major defoliator of potato, causing significant yield losses and pesticide use in North America, Europe, and Asia. CPB is difficult to control, in part, because it rapidly evolves resistance to insecticides (Alyokhin *et al*. 2008, Roush and Tingey 1992). Population differences in sensitivity to new insecticides may impact the effectiveness of those insecticides and the eventual evolution of CPB resistance. Identification of regional differences in baseline sensitivity to novel insecticides, when present, and the genetic mechanisms of those differences, will facilitate the development and evaluation of resistance management protocols for new insecticides.

When new insecticides employing novel modes of action are first tested on target pests, those pest populations will demonstrate variation in sensitivity among individuals. That variation may be due to genetic differences, environmental factors, or random differences in exposure among the tested insects. Good bioassay methods will minimize the influence of environmental and stochastic sources of variation while revealing the genetic sources. When populations of the target pest from different geographic locations are tested, they may also differ, possibly reflecting differences among the populations’ historical exposure to toxins or stochastic genetic differences among populations. These two measures of variation in sensitivity to the toxin, within and among populations, provide a basis for establishing application rates and are preliminary indicators of the risks of resistance evolution (ffrench-Constant and Roush 1990). It is not unusual to observe small but repeatable differences in sensitivity to a novel insecticide among geographic locations (Olson *et al*. 2000, Hitchner *et al*. 2012). Those differences are expected to occur randomly, with populations at different locations having equal probabilities of being more or less sensitive to a novel insecticide. This is because genetic differences among populations would be random with respect to the chemistry of novel insecticides acting on different target sites within the pest insect. Much more interesting, however, is when individuals from one population show relatively low or high sensitivity to a diverse array of novel insecticides, despite their differing modes of action. This could indicate that those populations have enhanced detoxification and excretion for that broad array of chemicals or that they are generally more robust or sensitive to toxin insult.

Geographic populations of CPB in the U.S. differ in their history of colonization. The beetles originated in the plains of the U.S., and, after a host shift onto potato, swept eastward colonizing the Midwest, Great Lakes and Atlantic coast regions in order (Tower 1906, Riley 1876, Izzo *et al*. 2018), and then later spread to the northwestern states of Washington, Oregon and Idaho, likely from already colonized potato growing regions in the East or Midwest (Tower 1906, Izzo *et al*. 2018). Because the potato agroecosystems in the Great Lakes region and East differ in several important respects from those of the Northwest (but not whether or not insecticides are applied), it is likely that CPB populations in those regions may differ in important ways (Clements *et al*. 2018, Crossley *et al*. 2019). It is not expected however that baseline sensitivity of CPB to insecticides with a diverse array of modes of action would systematically differ among these regions, as potato in all of these regions is treated with similar insecticides to control CPB and other insects (USDA NASS 2017).

Here, we analyzed an aggregate data set of independent bioassay results obtained over 15 years to ask the following question: are there geographic differences in CPB baseline sensitivity to novel insecticides? The bioassays were done before or soon after the first commercial use of new insecticides to assess the initial sensitivity of CPB to them. Because there was minimal previous exposure to the insecticides (in most cases none), we expect that sensitivity differences observed among individuals or populations from different locations are not caused by direct responses to selection imposed by the insecticides tested. To address possible molecular mechanisms underlying differences among populations, we performed an RNA-Seq analysis of expression differences between 2 populations that systematically differed in sensitivity to novel insecticides. By comparing gene expression of individuals from these two populations that are both unexposed and exposed to the insecticide imidacloprid, we seek to uncover candidate gene regulatory mechanisms that underlie physiological responses to insecticides in this insect.

## Material and Methods

### CPB populations

Between 1994 – 2009, CPB populations from a geographically wide range of potato farms across the USA and Canada were tested for sensitivity to new insecticides (Figure 1). In most cases, bioassays were performed before or soon after registration of the insecticide had occurred and therefore before its widespread commercial use in potato. Table 1 shows the number of farms sampled each year and tested with each insecticide.

**Table 1.** Numbers of populations of Colorado potato beetle from each of four geographic regions within the USA and Canada tested for each of six insecticides in this study.

| Regions | Insecticides |  |  |  |  |  |
| --- | --- | --- | --- | --- | --- | --- |
|  | Abamectin | Chlorantraniliprole | Cyantraniliprole | Imidacloprid | Metaflumizone | Spinosad |
| Great Lakes | 7 | 16 | 5 | 16 | 10 | 9 |
| Mid-Atlantic | 7 | 18 | 8 | 61 | 5 | 10 |
| North East | 3 | 14 | 5 | 7 | 7 | 10 |
| North West | 3 | 6 | 4 | 8 | 3 | 5 |

**Figure 1.**
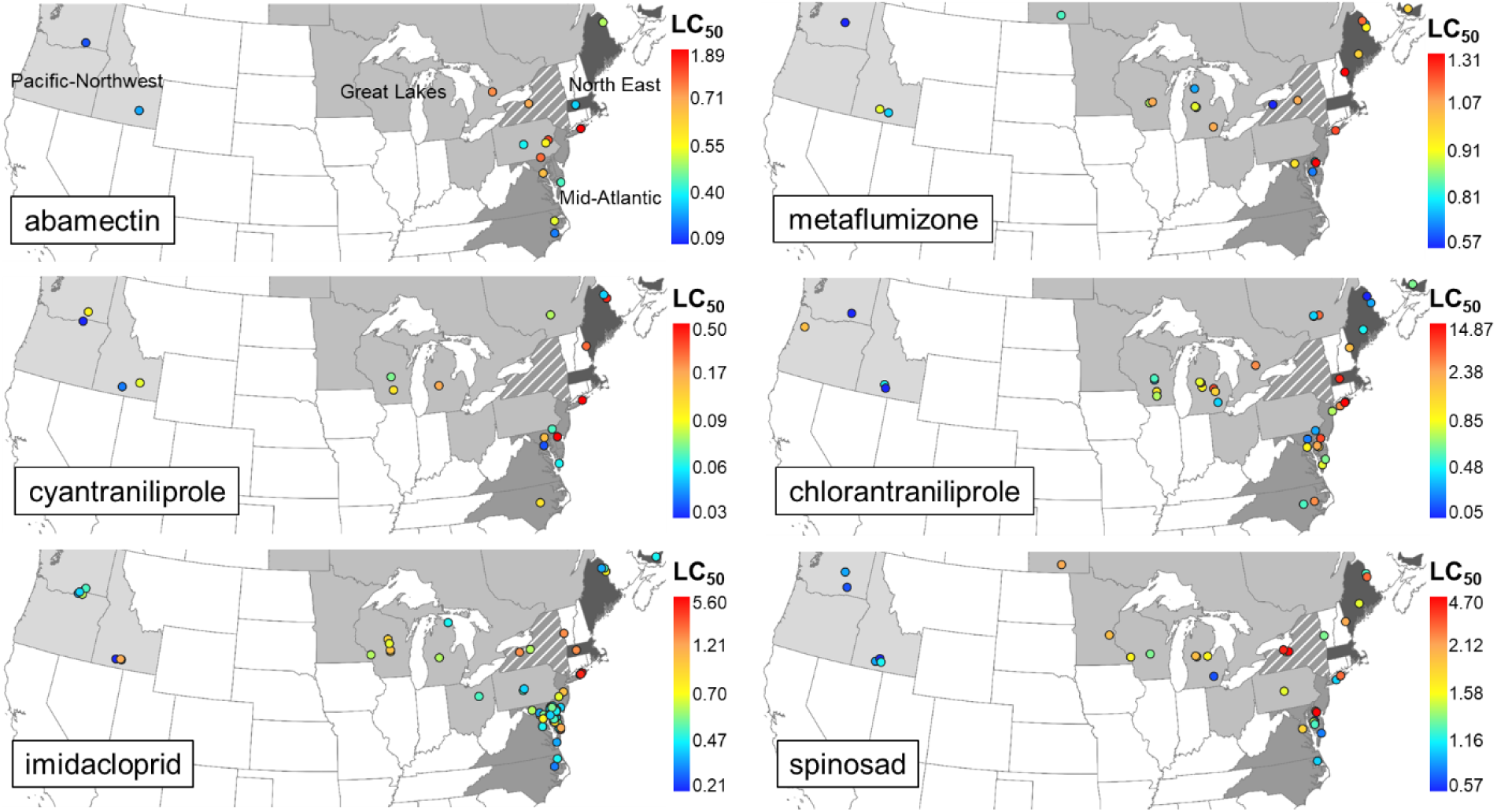
The location of CPB samples used to test effectiveness of new insecticides between 1994 – 2009. Circles indicate sample locations and are colored according to LC_50_. States are shaded to indicate their regional membership. New York is striped because samples from that state are in the Great Lakes or the Mid-Atlantic regions depending on their location.

Collections of 200-300 adult beetles from each farm were shipped by cooperators to the University of Maryland. Each population was segregated and reared in screened cages over potato plants in the field or in a laboratory environmental room. Egg masses attached to leaf discs were collected several times weekly and either stored at 12°C until larvae were needed or hatched immediately for bioassays. For all bioassay tests, neonate larvae were transferred to fresh potato leaves in Petri dishes and allowed to feed for 5 hours before testing. For collections with limited numbers of adults, the population was reared through one generation and tests were performed on the F1 progeny.

### Insecticides

The insecticides tested included the active ingredients abamectin, imidacloprid, spinosad, chlorantraniliprole, cyantraniliprole, and metaflumizone. Abamectin (MK-936) is a mixture of avermectins containing >80% avermectin B1a and <20% avermectin B1b and was obtained from Merck & Co. The product used was a 15% emulsifiable concentrate that was serially diluted in water and triton-X for bioassay. EPA residue tolerances were established for abamectin use in potatoes in 1999 and therefore not used in the U.S.A before then. We performed bioassays of CPB sensitivity to abamectin with 20 populations in 1994. Imidacloprid is a neonicotinoid insecticide that was tested as the formulated product (Admire 2 F, 21% [AI]]; Bayer, Kansas City, MO). The residue tolerance was established for imidacloprid in 1994 and we performed bioassays of imidacloprid on 34 populations in 1995, 50 populations in 1996 and 8 in 1999. Spinosad is a fermentation product of *Saccharopolyspora spinose*, containing two spinosyns (A and D). The formulated product, Spintor 2SC (Dow AgroSciences), was used in bioassays. Residue tolerances for spinosad were established in 1999 and we bioassayed 8 populations in 1999 and 16 in 2006. Chlorantraniliprole and cyantraniliprole are the active ingredients of two anthranilic diamide insecticides (rynaxypyr and cyazypyr, respectively (DuPont Agricultural Products) that act at ryanodine receptors of insects. The formulated products containing these insecticides were used for bioassays. Residue tolerances for chlorantraniliprole were set in 2008 and we bioassayed 16, 17, and 15 populations in 2006, 2007 and 2008 respectively. Twelve and 10 populations of cyantraniliprole were tested in 2008 and 2009 respectively; the residue tolerances were established in 2014. Metaflumizone is a semicarbazone insecticide that is a voltage-dependent sodium channel blocker. A formulated product of metaflumizone (BAS 320, a gift of BASF) was diluted with distilled water before use in bioassays. Residue tolerances have not yet been set for use of metaflumizone on potato so there has been no legal use of that compound on potatoes in the U.S.A. We bioassayed 25 populations of CPB in 2006 with that insecticide. All insecticides were diluted in distilled water (unless noted) before use in bioassays.

### Bioassay Methods

Each population was tested for susceptibility by exposing neonates (<24h post-hatch) to 6-8 concentrations of the test insecticides incorporated into a potato leaf-based agar diet and held in a growth chamber at 25°C (as described by Olson *et al*. 2000, Hitchner *et al*. 2012). For each insecticide, at least four replicate bioassays were performed with a minimum of 40 larvae tested per concentration. Mortality was assessed at 24 h intervals (most often 48 or 96 h depending on insecticide). Tests were repeated if control mortality exceeds 20% or if the concentration series did not produce an appropriate range of mortality around the 50% lethal level. In 6 of the 8 years, a laboratory population of CPB maintained by the New Jersey Department of Agriculture was tested along with populations collected from farms. This population was maintained without addition of individuals or exposure to pesticides to serve as a control population that would facilitate comparisons among bioassays. This control population was unavailable in 2008 and 2009. POLO-PC probit regression program (LeOra Software 1987) or Proc Probit in SAS (SAS Institute, *SAS User’s Guide*, ed. by SAS Institute, Cary N Cary, NC (1997)) were used to model the concentration-mortality responses to estimate the response slope, 50% lethal concentrations, and 95% confidence limits.

### Bioassay Analysis

Data from 8 years of the baseline bioassays were aggregated into a single data set. For each bioassay, year collected, chemical tested, farm location, and LC_50_ were recorded. When the NJ control population was tested in the same bioassay, a susceptibility ratio was calculated by dividing the LC_50_ of the farm population by the LC_50_ of the susceptible control. In order to compare bioassay results by region, farm populations were categorized into regional samples: MA = Mid-Atlantic Region (NJ, DE, MD, VA, NC), NE = North East (Eastern NY including Long Island, ME, MA, PEI), GL = Great Lakes (Western NY, QC, ON, PA, OH, MN, MI, MB, WI), and NW = Northwest (ID, OR, WA). Table 1 lists the number of CPB populations from each geographic region in the US and Canada assayed for each insecticide.

Values of LC_50_ and the susceptibility ratio were used to test for regional differences in sensitivity to each insecticide exposure. We statistically modelled the effect of each insecticide and tested for regional effects on LC_50_ and susceptibility ratio values of those bioassays that included the NJ control population.

Main effects of region, insecticide and their interaction were modeled as fixed factors using mixed model ANOVA (SAS Institute 1997), with bioassay year treated as a random effect. Before analysis, we inspected residual plots and performed the Shapiro-Wilk’s W test to examine for data normality and homogeneity of variances. Appropriate transformations or variance groupings were applied if necessary to satisfy the assumptions of ANOVA. Significant effects among means were separated using Tukey’s adjustment for pairwise comparisons (*P* ≤ 0.05).

## Transcriptional Differences of Regional Samples Materials and Methods

### Source of Insects

Adult CPB were collected in Hermiston, OR and Riverhead, NY (Long Island) and shipped to the University of Maryland in 2014. Beetles were contained in mesh cages on greenhouse grown potatoes. Leaves containing egg masses were collected and held at 25°C until hatching.

### Exposure treatment and extraction of RNA

First instar larvae were placed onto discs of potato leaf tissue that had been surface-treated with H_2_O or with 1.25 ppm Imidacloprid (Admire, Bayer) for 4 h before RNA extraction. This dosage of imidacloprid caused 8 and 0% mortality after 24 h to Oregon and New York larvae respectively. RNA from three replicate pools of larvae from each population (Oregon and New York) was extracted and used to create three replicate sequencing libraries for each location. Total RNA was extracted using a combined Trizol (Invitrogen) followed by RNAeasy (Qiagen) protocol. Each pool from New York beetles contains one larva from each of 12 sibships (same dam) and from 6-9 sibships of Oregon beetles.

### Sequencing and analysis of RNA-Seq data

Library construction and Illumina sequencing (150 bp paired-end) was performed by the University of Maryland Core Genomics facility. Reads from each sample were aligned to the CPB reference genome (PRJNA171749) using HISAT2 (Kim et al. 2015). Read counts were generated per sample per gene in the CPB official gene set (OGSv1.1) (Schoville et al. 2018) using the functions *makeTxDbFromGFF, transcriptsBy*, and *summarizeOverlaps* available in the R packages ‘GenomicAlignments’ and ‘GenomicFeatures’ (Lawrence et al. 2013). Overall gene expression patterns were visualized using Principle Components Analysis (PCA) on rlog-transformed read counts per gene per sample (Love *et al*. 2014). Using the resulting counts, we evaluated evidence for differential gene expression for each geographic or treatment groups using DESeq2 (Love *et al*. 2014). We retained genes for which differences in read counts were > 2-fold and significant at the α = 5% level. False discovery rates were controlled at the 1% level, using a Benjamini-Hochberg correction (Benjamini and Hochberg 1995). We retrieved annotations for genes for which manual curation had not been completed in the CPB OGS using Blast2GO (Conesa *et al*. 2005) annotations generated by Crossley *et al*. (2017). R code used for read counting and differential expression analysis is available in Supp. File 1.

## Results

### Regional differences in baseline bioassays

Figure 2 shows the results of a series of bioassays conducted between 1994 – 2009. In all cases, the median LC_50_ of the NW samples was lower than those of the NE, GL, or MA samples. Perhaps more importantly, for all insecticides, populations with the highest baseline tolerance were either Mid-Atlantic or Northeastern.

**Figure 2.**
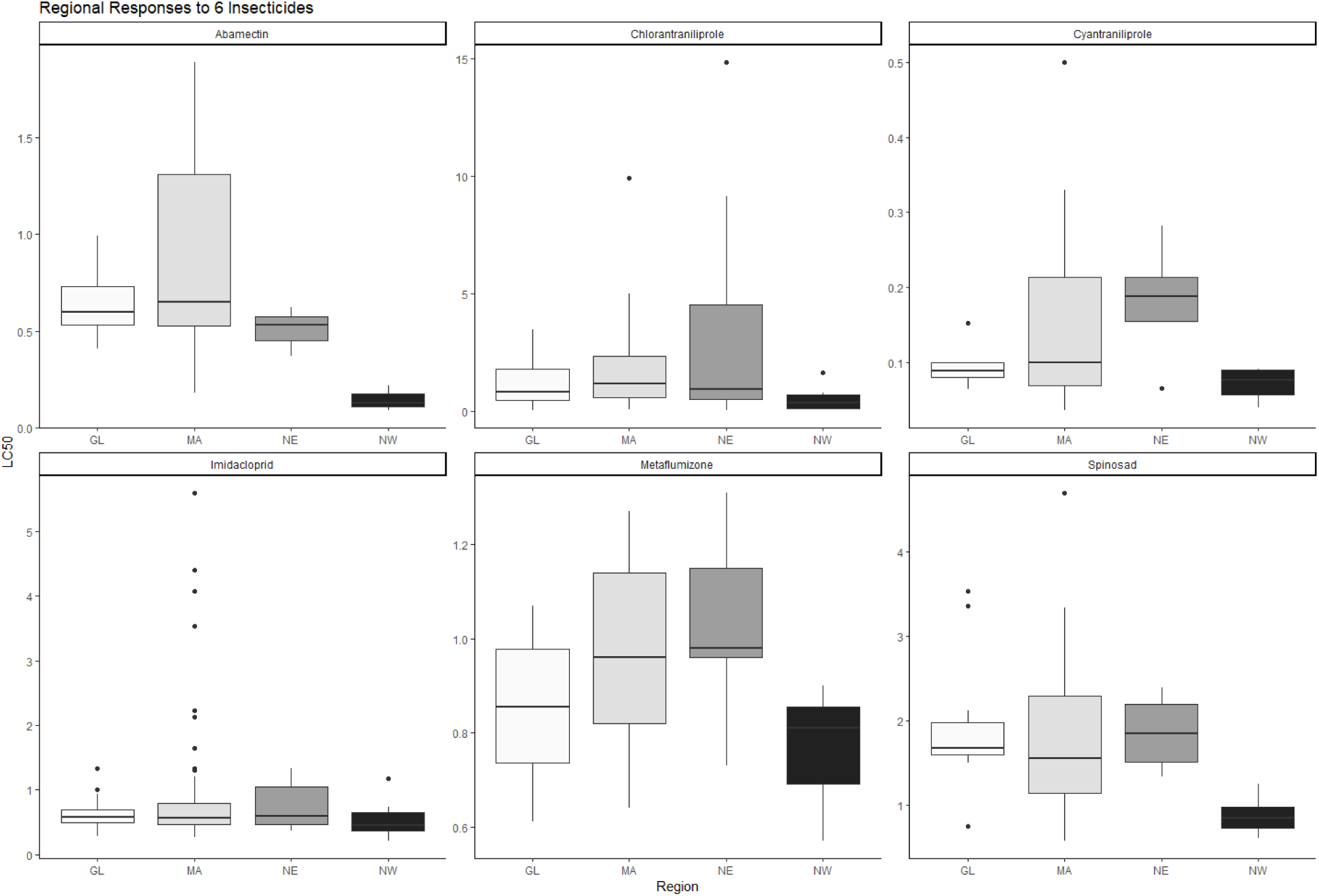
Regional bioassay responses of CPB to six insecticides. The box contains the middle 50% of observation (with cutoffs at 25^th^ and 75^th^ percentiles). The line within each box is the median LC_50_. The upper and lower whiskers extend from the box edges to the largest or smallest value (respectively) no further than 1.5x the distance between the first and third quartiles. Observations beyond that are indicated by a solid circle. Regions: GL = Great Lakes (Western NY, QC, ON, PA, OH, MN, MI, MB, WI), MA = Mid-Atlantic Region (NJ, DE, MD, VA, NC), NE = North East (Eastern NY including Long Island, ME, MA, PEI), and NW = Northwest (ID, OR, WA).

Analysis of bioassay data revealed similar significant differences among regions (*P* < 0.0001, 3 df) and chemicals (*P* < 0.0001, 5 df) using either LC_50_ or the relative sensitivity ratio as response variables (Table 2). The interactions of ‘Region’ and ‘Chemical’ were not significant for either LC_50_ or the relative sensitivity ratio, indicating that the relative performance of each region did not differ across the different insecticides. Adjusted Tukey’s tests of differences among regions revealed significant differences between NW and the other three regions (P< 0.001), but not among the remaining three regions.

**Table 2.** Analysis of variance of LC_50_ and Sensitivity ratios.

| **LC<sub>50</sub>** |  |  |  |  |
| --- | --- | --- | --- | --- |
| Effect | Num DF | Denom DF | F Value | Pr > F |
| Chemical | 5 | 31.1 | 14.2 | < 0.0001 |
| Region | 3 | 92.7 | 14.14 | < 0.0001 |
| Chemical * Region | 15 | 88.9 | 1.51 | 0.1185 |
| **Sensitivity ratio** |  |  |  |  |
| Effect | Num DF | Denom DF | F Value | Pr > F |
| Chemical | 4 | 20.7 | 37.10 | < 0.0001 |
| Region | 3 | 78.7 | 11.91 | < 0.0001 |
| Chemical * Region | 12 | 79.9 | 1.6 | 0.1083 |

### RNA-Seq comparison of NY and OR baseline expression

Illumina sequencing generated a total of 232 million paired-end reads, with 15.9 to 22.9 million reads obtained per sample. Alignment rates to the reference genome ranged from 85% to 91%. Overall gene expression patterns were more variable between CPB populations than between imidacloprid-treated and untreated beetles within populations (Fig. 3). In fact, no genes were significantly differentially expressed between imidacloprid-treated and untreated beetles in either population. This may have resulted from a relatively low exposure dosage and/or short time between the treatment and RNA extraction. Comparison of gene counts in untreated beetles between regions, however, revealed 28 significantly up-regulated and 44 down-regulated genes in samples from the New York relative to the Oregon population (Fig. 3a). A large proportion of genes with increased expression were associated with metabolic detoxification of insecticides, with differences in gene counts ranging from 3-to 11-fold between populations; 12 up-regulated genes belong to the cytochrome P450 gene family and one esterase (Table S1, Fig. 3). Two target-sites of older insecticides also showed increased expression in the eastern population (3.5-fold each): acetylcholine esterase, the target of organophosphates and carbamates (IRAC mode of action Groups 1A and 1B) (IRAC MOA Classification 2019), and the gamma-aminobutyric acid receptor, target of organochlorines (IRAC MOA Group 2). Genes with reduced expression in the eastern population comprised a large number of genes encoding cuticular proteins (16 out of 44), three gustatory receptors, a protease, a glutathione-*S*-transferase (associated with metabolic detoxification), and a transient receptor potential cation channel (target of IRAC MOA Group 9B, pyridine azomethine derivatives).

**Figure 3.**
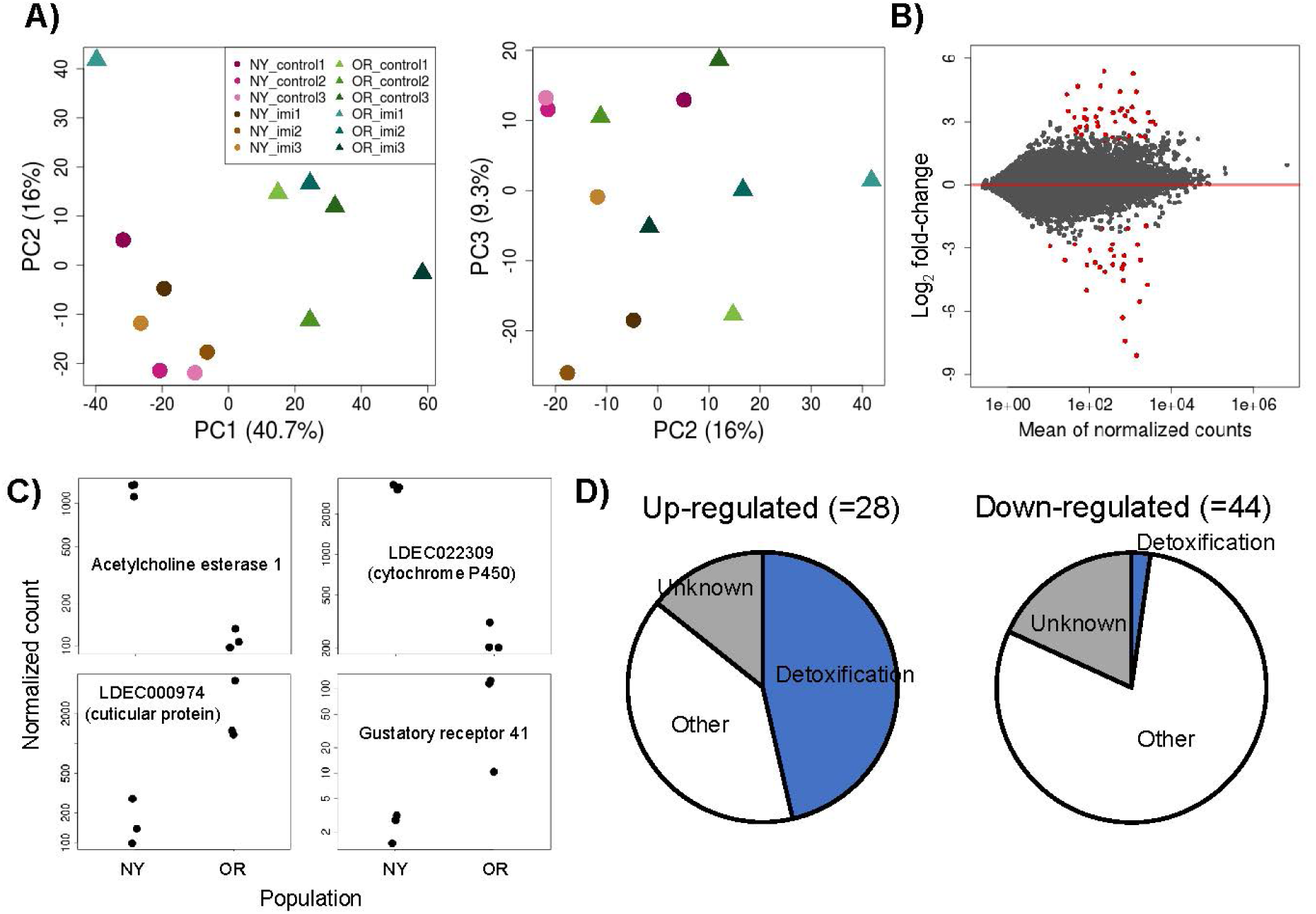
A) Principal Components Analysis of rlog-transformed read counts per *L*. *decemlineata* gene. B) MA plot depicting the relationship between mean transcript counts (mean of normalized counts) and the differences in transcript counts (log_2_fold-change) between untreated New York and Oregon CPB. D) Pie charts depicting proportional representation of gene families among genes constitutively up- and down-regulated in the New York relative to Oregon CPB population samples. C) Example difference in read counts of acetylcholine esterase 1, LDEC022309 (a cytochrome P450), LDEC000974 (cuticular protein), and gustatory receptor 41, in New York and Oregon CPB population samples.

## Discussion

Analysis of a multi-year, multi-insecticide set of bioassays provides clear evidence of regional differences in CPB sensitivity to a diverse array of novel insecticides. The Northwestern populations from Oregon and Washington were more sensitive to all compounds and the Northeastern, Mid-Atlantic and Great Lakes populations were less sensitive. These regional differences reflect pre-selection sensitivity to a novel toxin, because they are not the result of direct selection for increased tolerance to the insecticides that were tested nor to other insecticides from the same class. This does not mean that the populations have not experienced selection by insecticides, as they almost certainly have in the more than 100 years of chemical control targeting this pest. At the time of bioassay, however, the populations had not faced these novel chemicals nor their unique modes of action. Although these bioassays reveal significant differences in the phenotypes of CPB from different regions, the tested insecticides were initially effective in the field at all locations. It is also important to note that although CPB from the Northeastern, Mid-Atlantic and Great Lakes regions are less sensitive on average than those in the Northwest, there are samples from those eastern regions that were quite sensitive, though none from the Northwest that were among the most tolerant.

The target sites of these six insecticides, as currently understood, may not be unique, but they represent at least five different modes of action: glutamate gated chloride channel (abamectin, IRAC MOA Group 6), ryanodine receptor (cyantraniliprole and chlorantraniliprole, IRAC MOA Group 28), nicotinic acetylcholine receptors (imidacloprid and spinosad, IRAC MOA Groups 4A and 5, respectively), and sodium channels (metaflumizone, IRAC MOA Group 22B) (IRAC MOA Team 2019). With the exception of chlorantraniliprole, they are also the first members of the chemical class to be considered or approved for use in potato. Insect sensitivity to cyantraniliprole and chlorantraniliprole may reasonably be expected to be positively correlated because they both target the ryanodine receptors of insects and therefore may share target sites. Imidacloprid and spinosad both target the nicotinic acetylcholine receptors, but in different ways, and their correlated affinity to target sites remains to be determined (Sparks and Nauen 2015, Mota-Sanchez *et al*. 2006). Because the populations tested here were naïve with respect to the tested insecticides, we suspect that regional variation in sensitivity is due to differences in metabolic, excretion, and exclusion mechanisms that may be adaptations to generalized chemical insult resulting from years of exposure to other insecticides or fungicides, and not to target site insensitivities to the tested compounds (Clements *et al*. 2018).

The RNA-Seq analysis identified a number of genes coding for detoxification-related products that are constitutively up regulated in the New York population, relative to the Oregon population, supporting the hypothesis that the regional differences in baseline sensitivity may be due to increased expression of genes for generalized metabolism and excretion of chemical toxins. Population level differences in the regulation of detoxification genes are correlated with increased tolerance to insecticides in several pest insects, including CPB (Kalsi and Palli 2017, Bass *et al*. 2014). The numerous CYP genes and the esterase gene with increased expression in the New York population are notable, as is the increased expression of two target sites (ACE and GABA) of legacy organophosphate and organochlorine insecticides. We also note the decreased expression of multiple cuticle protein genes in the New York population. Increased expression of cuticular proteins has been associated with increased tolerance of insecticide exposure because of reduced penetration of the toxin (Balabanidou et al. 2018). Here, the more tolerant population (NY) has reduced expression of several cuticular proteins, suggesting that another explanation is required. While these detoxification-related genes may be independently regulated and constitutively increased following a series of independent changes, it is also possible that they are co-regulated by a few, or perhaps a single, upstream regulatory process such as the CncC pathway thought to underlie an expression-based resistance to imidacloprid in this insect (Gaddelapati et al. 2018). Although it is beyond the scope of this paper to identify the causes of this regional difference, it seems most likely that it is a result of previous pesticide exposure histories, though the details of that exposure remain unknown. Further, it is possible that the expression levels of these generalized detoxification-related genes may serve as an indicator of the potential for CPB adaptation to novel insecticides, especially those that are the substrates of these enzymes. Because the insects used in the RNA-Seq were sampled in 2014 after all of the bioassays had occurred, we do not know when the constitutive increased expression of detoxification genes first occurred and we cannot say with certainty that these changes are the cause of the regional differences in sensitivity observed.

In addition to regional differences in expression of genes involved in metabolic detoxification, we observed relatively higher expression of a suite of cuticle protein genes in Oregon CPB. The reduced expression in New York CPB is not consistent with research finding a positive association between cuticle protein gene expression and reduced penetration of insecticides (Balabanidou et al. 2018). However, high expression of cuticle protein genes in eastern Oregon, which receives 40-fold less annual precipitation than New York does, is consistent with greater selection for desiccation resistance among Oregon CPB. Decreased baseline expression of genes involved in metabolism is also consistent with selection for reduced respiration (and consequent water loss) in a drier climate (Hoffmann and Harshman 1999). Thus, an alternative explanation to regional differences in baseline sensitivity to insecticides is that the drier climate of eastern Oregon imposes constraints on baseline expression of genes involved in metabolism, limiting eastern CPB’s propensity to rapidly evolve resistance to insecticides through generalized metabolism and excretion.

Several additional explanations for the regional bioassay and expression differences are unlikely. First, although some estimates of genome-wide genetic diversity indicate that Northwestern populations have reduced overall genetic variation compared to eastern populations, these differences would not result in different levels of baseline tolerance to a new insecticide, though they may contribute to differences in the rate of adaptation of CPB to an insecticide. When genetic variation in the coding regions of candidate resistance-causing genes is measured, there is no reduction in diversity observed in populations sampled from the NW (S. Schoville, unpublished data). Second, we also do not expect the differences in detoxification-related gene transcript levels to be due to differing field exposures of the tested individuals, because we used the unexposed offspring of collected individuals for RNA-Seq analysis. Finally, the observed regional differences in gene expression, which agree with regional differences in sensitivity to six insecticides, are unlikely to be due to stochastic differences in the expression level of detoxification genes. The combination of similar relative levels of sensitivity across several insecticide chemistries and the increased expression of a suite of detox genes in one (the less sensitive) but not the other region strongly suggest that the pattern is not random. Finally, because the RNA-Seq libraries were constructed from pooled RNA of offspring of 10-12 field-sampled females, it is unlikely that the observed patterns are due to sampling of rare genotypes. Additional work is needed to confirm that the regional differences are indeed genetic, and to identify the sources of those genetic differences.

Recent tests of imidacloprid resistance show that Northwestern populations remain susceptible, despite more than 20 years of exposure, whereas high levels of tolerance have evolved in the Great Lakes, Northeastern and Mid-Atlantic regions (Crossley et al. 2018, Alyokhin et al. 2015, Szendrei et al. 2012). This observation of persistent regional differences in sensitivity to novel insecticides is important in several ways. First, it suggests that the initial effectiveness of novel insecticides and the application concentrations should be determined using CPB from both the more and less tolerant regions, with field application rates potentially differing geographically to reflect baseline differences. Second, the initial tolerance of populations to a novel toxin may play a role in the rate or likelihood of the evolution of resistance to that toxin. The many cases of insecticide resistance in CPB have been first observed in the eastern half of the USA (Alyokhin *et al*. 2015, Forgash 1985). Some of these cases of resistance may have benefitted from greater baseline tolerance of eastern populations to the insecticide. If this is the case, we may be able to incorporate baseline sensitivity of populations to a novel insecticide in the resistance risk analysis, focusing monitoring efforts on those populations with the higher initial tolerance even when all populations are equally well-controlled in the field. Interestingly, this suggests a mechanism for the “Pesticide Treadmill” in which pest populations that have evolved resistance to chemical control rapidly evolve resistance to subsequent chemically distinctive toxins. In addition to the loss of natural enemies due to insecticide overuse, populations may evolve higher baseline tolerances to a diverse array of toxic compounds through the increased expression of generalized detoxification mechanisms, leading to more-rapid evolution of resistance to the next insecticide. This is similar to the case with *Myzus persicae* in which populations that show initial tolerance to pyrethroids or neonicotinoids because of enhanced detoxification were also the first to evolve field-relevant levels of resistance to those insecticides (Bass *et al*. 2014). If this is the case, then we may need to assess the risks of cross-resistance more broadly than considering only target site similarity (Sparks and Nauen, 2015). If an insecticide selects for a more generalized response through increased expression of metabolic, excretion, or absorption related genes, it may increase the risks of field-level resistance in a subsequent insecticide even when the insecticides do not share target sites. Finally, the observed differences in baseline tolerance to novel insecticides could cause responses to selection by those insecticides to differ, leading to geographic differences in the genetics of tolerance to those toxins.

## Acknowledgments

We are grateful to Silvia Rondon, who provided beetles for the RNA-Seq analyses. S.D.S. and M.S.C. were supported by Hatch Act formula funds (#WIS02004) and the USDA AFRI-ELI Graduate Fellowship (#2018-67011-28058). D.J.H. was supported by a USDA-ARS Potato Production and Disease Research Grant and by the National Socio-Environmental Synthesis Center (SESYNC) under funding received from the National Science Foundation DBI-1052875.

