## Supplemental Table 1 for "Regional differences in gene regulation may underlie patterns of sensitivity to novel insecticides in Colorado potato beetle"

**Table S1.** Differentially expressed genes in the Riverhead, New York and Hermison, Oregon populations of Colorado potato beetle, *Leptinotarsa decemlineata.* Gene identifications based on the *L. decemlineata* official gene set (OGSv1.1) (Schoville et al. 2018) and annotations reported in Crossley *et al.* (2017).

| **Control OR vs. LI** |  | **(-) = upregulated in LI; (+) = upregulated in OR** |  |
| --- | --- | --- | --- |
| **Gene** | **Gene Identification** | **log2foldchange ± SEM** | **Adjusted p-value** |
| LDEC023780 | cytochrome p450 | -10.9±1.1 | 6.66E-18 |
| LDEC020527 | cytochrome P450 9e2-like | -9.7±0.6 | 6.80E-51 |
| LDEC022531 | esterase | -7.8±1.1 | 1.13E-07 |
| LDEC020528 | Gag-pol poly | -7.5±0.5 | 2.25E-35 |
| LDEC021674 | Transposon Ty3-G Gag-Pol poly | -6.7±1.4 | 0.006639429 |
| LDEC020421 | cytochrome P450 | -6.1±0.3 | 5.93E-46 |
| LDEC010984 | polycystic kidney disease protein 1-like 2 | -5.2±0.7 | 3.84E-07 |
| LDEC017299 | PREDICTED: uncharacterized protein LOC107399201 | -5.1±0.8 | 0.000228957 |
| LDEC020419 | ---NA--- | -4.9±0.2 | 1.28E-79 |
| LDEC013538 | cytochrome p450 6bq10 | -4.9±0.5 | 2.07E-13 |
| LDEC021334 | cytochrome P450 | -4.8±0.3 | 2.51E-41 |
| LDEC023239 | cytochrome p450 | -4.3±0.4 | 1.81E-17 |
| LDEC022491 | PREDICTED: uncharacterized protein LOC106707885 | -4.3±0.4 | 3.67E-13 |
| LDEC001159 | neurexin-1 isoform X2 | -4.2±0.7 | 0.000308724 |
| LDEC022856 | cytochrome p450 | -4.2±0.4 | 2.22E-15 |
| LDEC021333 | cytochrome P450 9e2-like | -4.1±0.2 | 7.66E-50 |
| LDEC001160 | hypothetical protein TcasGA2_TC034204 | -4.0±0.7 | 0.00589811 |
| LDEC024396 | cytochrome P450 9e2 [Tribolium castaneum] | -4.0±0.2 | 6.41E-30 |
| LDEC022309 | cytochrome p450 | -3.8±0.2 | 6.83E-28 |
| LDEC018533 | cytochrome P450 | -3.6±0.3 | 9.55E-19 |
| LDEC005342 | gamma-aminobutyric acid receptor alpha-like | -3.6±0.4 | 1.03E-06 |
| Acetylcholine esterase 1 | NA | -3.5±0.2 | 2.38E-25 |
| LDEC022391 | ---NA--- | -3.4±0.5 | 0.00045195 |
| LDEC014763 | sodium-coupled monocarboxylate transporter 1 | -3.2±0.4 | 1.84E-05 |
| LDEC006940 | cytochrome P450 9e2-like | -2.9±0.2 | 5.04E-25 |
| LDEC003363 | p protein | -2.2±0.3 | 0.004448206 |
| LDEC022100 | chymotrypsin inhibitor-like | -2.1±0.2 | 0.000457231 |
| LDEC016889 | prostatic acid phosphatase | -2.0±0.2 | 0.000246724 |
| LDEC000957 | isoform b | 2.2±0.3 | 0.004580215 |
| LDEC017909 | carbohydrate-binding protein | 2.3±0.3 | 0.001300095 |
| LDEC018083 | ---NA--- | 2.4±0.3 | 0.000218015 |
| LDEC018084 | collagen alpha-1 chain | 2.4±0.3 | 0.000261101 |
| LDEC013059 | hypothetical protein TcasGA2_TC001141 | 2.5±0.4 | 0.009317022 |
| LDEC007014 | spermine oxidase | 2.6±0.3 | 0.001023905 |
| LDEC023323 | glutathione s-transferase isoform d | 2.7±0.4 | 0.003896166 |
| LDEC009472 | ---NA--- | 2.7±0.4 | 0.002130325 |
| LDEC013670 | transient receptor potential cation channel subfamily v member 6 | 3.0±0.5 | 0.004827524 |
| LDEC015141 | titin isoform x2 | 3.1±0.4 | 7.60E-06 |
| LDEC005840 | serine ase stubble | 3.1±0.5 | 0.002189917 |
| LDEC015111 | pupal cuticle protein edg-84a-like protein | 3.4±0.5 | 0.002130325 |
| LDEC021147 | gag-pol poly | 3.6±0.6 | 0.005293498 |
| LDEC008124 | cysteine sulfinic acid decarboxylase | 3.7±0.5 | 0.000144072 |
| LDEC000974 | cuticular protein 4 | 3.7±0.5 | 0.00017818 |
| LDEC013984 | tpa: cuticle protein | 3.8±0.5 | 2.42E-05 |
| LDEC001805 | flexible cuticle protein 12-like | 3.9±0.6 | 0.000308724 |
| LDEC005441 | rna polymerase ii degradation factor 1 | 3.9±0.6 | 0.000557475 |
| LDEC015104 | larval cuticle protein a2b-like | 4.0±0.7 | 0.002130325 |
| LDEC023912 | larval cuticle protein a2b-like | 4.0±0.6 | 0.000438766 |
| LDEC010635 | protein wnt-11b-2-like | 4.1±0.6 | 1.63E-05 |
| LDEC016285 | rna-directed dna polymerase from mobile element jockey-like | 4.1±0.6 | 5.09E-05 |
| LDEC001804 | flexible cuticle protein 12-like | 4.2±0.5 | 2.15E-08 |
| LDEC012326 | endocuticle structural glycoprotein bd-8-like | 4.2±0.5 | 6.84E-08 |
| LDEC016286 | cuticle protein 76-like | 4.3±0.7 | 0.000308724 |
| LDEC016822 | lish domain-containing | 4.5±0.5 | 6.85E-08 |
| LDEC001803 | cuticle protein | 4.5±0.6 | 3.32E-06 |
| LDEC024732 | larval pupal cuticle protein h1c-like | 4.5±0.6 | 3.17E-06 |
| LDEC015346 | anoctamin-10 isoform x2 | 4.6±0.6 | 8.96E-08 |
| LDEC015110 | cuticle protein 21-like | 4.8±0.6 | 6.85E-08 |
| LDEC004956 | hypothetical protein D910_12530 | 5.0±0.7 | 9.42E-07 |
| Gustatory receptor 41 | NA | 5.4±0.9 | 0.00045195 |
| LDEC019837 | ---NA--- | 5.4±0.5 | 9.55E-15 |
| LDEC015097 | larval cuticle protein a2b-like | 5.9±0.6 | 9.37E-13 |
| LDEC016824 | reverse transcriptase | 6.1±0.6 | 7.18E-14 |
| Gustatory receptor 42 | NA | 6.1±1.1 | 0.00045195 |
| LDEC015890 | ---NA--- | 6.2±0.9 | 1.00E-05 |
| LDEC016884 | gustatory receptor 125 | 6.5±0.8 | 1.37E-08 |
| LDEC016288 | cuticle protein 76-like | 6.5±0.5 | 1.00E-22 |
| LDEC016294 | coiled-coil domain-containing protein 142 | 6.6±1.1 | 9.50E-05 |
| LDEC015098 | larval cuticle protein a2b-like | 6.9±1.2 | 0.000557475 |
| LDEC017424 | ---NA--- | 7.2±0.6 | 1.05E-19 |
| LDEC015347 | ---NA--- | 7.8±1.0 | 2.15E-08 |
| LDEC009858 | ---NA--- | 8.1±1.3 | 2.42E-05 |
| **Treated OR vs. LI** |  |  |  |
| LDEC023780 | cytochrome p450 | -11.7±1.2 | 5.81E-16 |
| LDEC020527 | cytochrome P450 9e2-like | -9.2±0.6 | 5.67E-39 |
| LDEC020528 | Gag-pol poly | -6.3±0.6 | 1.81E-16 |
| LDEC021819 | salicyl alcohol oxidase paralog partial | -5.8±0.8 | 4.88E-07 |
| LDEC020421 | cytochrome P450 | -5.5±0.5 | 3.62E-14 |
| LDEC022531 | esterase | -5.2±0.6 | 1.98E-08 |
| LDEC017501 | hypothetical protein D910_09427 | -5.2±1.0 | 0.009694815 |
| LDEC023402 | 50s ribosome-binding partial | -5.2±0.8 | 3.90E-05 |
| LDEC016725 | ---NA--- | -5.0±0.7 | 8.44E-07 |
| LDEC022491 | PREDICTED: uncharacterized protein LOC106707885 | -4.6±0.5 | 1.79E-08 |
| LDEC021334 | cytochrome P450 | -4.5±0.4 | 5.30E-19 |
| LDEC016724 | reverse partial | -4.3±0.5 | 7.26E-08 |
| LDEC017037 | sodium channel protein nach | -4.2±0.7 | 0.000586888 |
| LDEC022856 | cytochrome p450 | -4.2±0.6 | 9.95E-06 |
| LDEC013538 | cytochrome p450 6bq10 | -4.2±0.6 | 1.24E-05 |
| LDEC020419 | ---NA--- | -4.1±0.3 | 8.99E-23 |
| LDEC023239 | cytochrome p450 | -4.0±0.6 | 7.15E-05 |
| LDEC021333 | cytochrome P450 9e2-like | -3.9±0.3 | 4.88E-15 |
| LDEC009289 | endopolygalacturonase | -3.8±0.7 | 0.008339451 |
| LDEC024396 | cytochrome P450 9e2 [Tribolium castaneum] | -3.7±0.3 | 9.88E-17 |
| Acetylcholine esterase 1 | NA | -3.5±0.2 | 3.61E-21 |
| LDEC001159 | neurexin-1 isoform X2 | -3.3±0.4 | 9.26E-07 |
| LDEC022309 | cytochrome p450 | -3.3±0.4 | 3.90E-05 |
| LDEC001160 | hypothetical protein TcasGA2_TC034204 | -3.1±0.5 | 0.006358154 |
| ABC transporter subfamily C member ABCC-76B | NA | -3.0±0.4 | 0.002132062 |
| LDEC018533 | cytochrome P450 | -2.8±0.3 | 4.41E-07 |
| LDEC019606 | MFS-type transporter | -2.6±0.4 | 0.007515605 |
| LDEC006940 | cytochrome P450 9e2-like | -2.5±0.3 | 0.002238276 |
| LDEC001646 | ---NA--- | 3.0±0.4 | 0.000241675 |
| LDEC003816 | PREDICTED: uncharacterized protein LOC100142369 | 3.1±0.5 | 0.003104567 |
| LDEC015141 | titin isoform x2 | 3.1±0.4 | 6.92E-06 |
| LDEC010635 | protein wnt-11b-2-like | 3.4±0.6 | 0.007068002 |
| LDEC013670 | transient receptor potential cation channel subfamily v member 6 | 3.7±0.4 | 3.08E-07 |
| LDEC011537 | ankyrin-2-like protein | 4.6±0.4 | 6.00E-16 |
| LDEC015890 | ---NA--- | 4.9±0.9 | 0.004003199 |
| LDEC004956 | hypothetical protein D910_12530 | 5.2±0.4 | 9.95E-26 |
| LDEC016884 | gustatory receptor 125 | 6.2±0.8 | 1.72E-08 |
| Gustatory receptor 42 | NA | 7.0±1.0 | 3.42E-07 |
| Gustatory receptor 41 | NA | 7.5±1.1 | 1.02E-05 |
| LDEC004957 | hypothetical protein YQE_11734, partial | 7.8±0.8 | 3.81E-14 |
